# Optimizing actin turnover in cell-like conditions

**DOI:** 10.64898/2026.02.02.703230

**Authors:** Fabina Binth Kandiyoth, Fabien Durbesson, Christine Hajjar, Xingbo Yang, Etienne Loiseau, Alphée Michelot

## Abstract

A dynamic actin cytoskeleton, characterized by rapid filament turnover, is essential for driving intracellular transport and cellular movement. Despite extensive study, the contributions of many actin-binding proteins (ABPs) that regulate the different steps of the actin turnover cycle (filament assembly, disassembly and recycling into polymerizable actin monomers) remain ill-defined. Here, we introduce novel sensitive in vitro assays to quantitatively assess how ABPs catalyze actin turnover. By accurately measuring nucleotide exchange dynamics and ATP consumption, these assays enable robust characterization of ABP activity across broad concentration ranges. Using these methods and modeling of these reactions, we systematically examined the contributions of five conserved regulators, both individually and in combination, and identified conditions that maximize turnover efficiency. We also determined that increasing F-actin concentration to cellular levels affects ABP activity. Finally, we demonstrated that rapid actin turnover is preserved during encapsulation in cell-sized vesicles using the cDICE method. Together, these advances provide versatile tools and new insights into actin cytoskeletal dynamics under physiologically relevant conditions.

## Introduction

Actin is a highly conserved and ubiquitous protein of the eukaryotic cell cytoskeleton. This ATPase continuously transitions between two forms, monomeric (or G-actin) and filamentous (or F-actin), in a process called ‘actin turnover’. The transition from the G-state to the F-state is a polymerization reaction, during which actin monomers bound to ATP nucleate new filaments or elongate pre-existing ones at their barbed ends. The return from the F-state to the G-state is called disassembly, a process that involves a cascade of complex events including filament fragmentation, bursting, and pointed-end depolymerization after ATP hydrolysis. Finally, completion of the turnover cycle requires efficient recycling of ADP-bound actin monomers into polymerizable ATP-bound ones [1,2].

Active polymerization is essential for actin to perform its motor function in numerous cellular processes [3]. However, actin alone is primarily stable in its filamentous form and capable only of very slow turnover. To overcome this limitation, cells express a variety of specialized actin-binding proteins (ABPs) that accelerate one or several of the main rate-limiting steps of the actin turnover cycle [4]. The expression of such a wide variety of proteins reflects a degree of redundancy that is crucial for essential cellular processes such as actin turnover and the fact that certain ABPs collaborate with each other.

The slow steps of actin turnover that need to be primarily catalyzed are actin disassembly and recycling [5]. During disassembly, single filaments are first dissociated from actin networks [6,7] and severed into shorter fragments by ADF/cofilin [8,9]. The cooperative binding of ADF/cofilin to F-actin [10] accelerates the release of inorganic phosphate resulting from ATP hydrolysis [11,12] and promotes filament severing at the interface between ADF/cofilin-decorated and bare sections of the filaments [13,14]. The disassembly of these fragments then occurs by rapid subunit-by-subunit depolymerization or by bursting, a fast second-scale process resembling the catastrophic disassembly of microtubules. Rapid depolymerization and bursting both involve ADF/cofilin and additional enzymes such as Aip1, Cyclase Associated Protein (CAP), and coronin [14–23]. It is not yet clear how depolymerization and bursting contribute individually to actin disassembly. One possibility is that bursting may not allow actin filaments to return completely to their monomeric state, and that rapid depolymerization is necessary to disassemble the small actin oligomers resulting from bursting.

Our understanding of these molecular mechanisms has advanced considerably in recent years, notably through techniques such as TIRF microscopy, which allow precise measurements of the effects of individual molecules on single filaments [24]. However, we still lack information on the collective actions of these proteins, on how their efficiencies compare, and on how they perform under cellular concentrations and volumes. This lack of information also limits the development of biomimetic assays in cell-like conditions [25]. To address these gaps, we have developed new sensitive, high-throughput, and easy-to-implement assays to characterize the collective behavior of actin-binding proteins (ABPs) involved in actin turnover. These assays enabled us to optimize experimental conditions for the reconstitution of rapid actin dynamics inside cell-like biomimetic systems, such as giant unilamellar vesicles (GUVs).

## Results

### Quantification of actin disassembly and recycling rates by release of fluorescent nucleotides

Previously, we identified a highly sensitive fluorescent ATP, N^6^-(6-amino)hexyl-ATP-ATTO-488 (referred to as ATP-488 in subsequent parts of the article), which binds to actin monomers (G-actin) [26]. ATP-488 does not prevent the actin polymerization reaction nor inhibits the binding of many actin regulators such as profilin or ADF/cofilin. Furthermore, the binding or dissociation of ATP-488 from G-actin can be detected by fluorescence anisotropy. Anisotropy signals are saturated when ATP-488 is bound to G-actin, and no longer increase as actin polymerizes. Notably, fluorescence anisotropy is not sensitive to photobleaching, and enables sensitive monitoring of nucleotide binding and release over hours.

We asked whether ATP-488 could be used to quantify and compare the efficiency of ABPs involved in actin disassembly and recycling, by measuring the release rate of the fluorescent nucleotides in solution from an F-actin pre-bound state (Fig. 1A). We developed a multi-step assay in which a fraction of ATP bound to G-actin is first exchanged for ATP-488 (Fig. 1A - step 1). Actin is then polymerized for two hours, a deliberately long duration chosen so that actin would reach a steady-state equilibrium [26]. At equilibrium, bound nucleotides are presumably hydrolyzed, and all subunits of the filament are bound to ADP or ADP-488 (Fig. 1A - step 2). While this dynamic equilibrium occurs, the combined rate of filament disassembly and recycling is quantified using fluorescence anisotropy by adding an excess of ATP (Fig. 1A - step 3). This excess ATP prevents any reassociation of ADP-488 with G- or F-actin once it has been released in solution.

**Figure 1.**
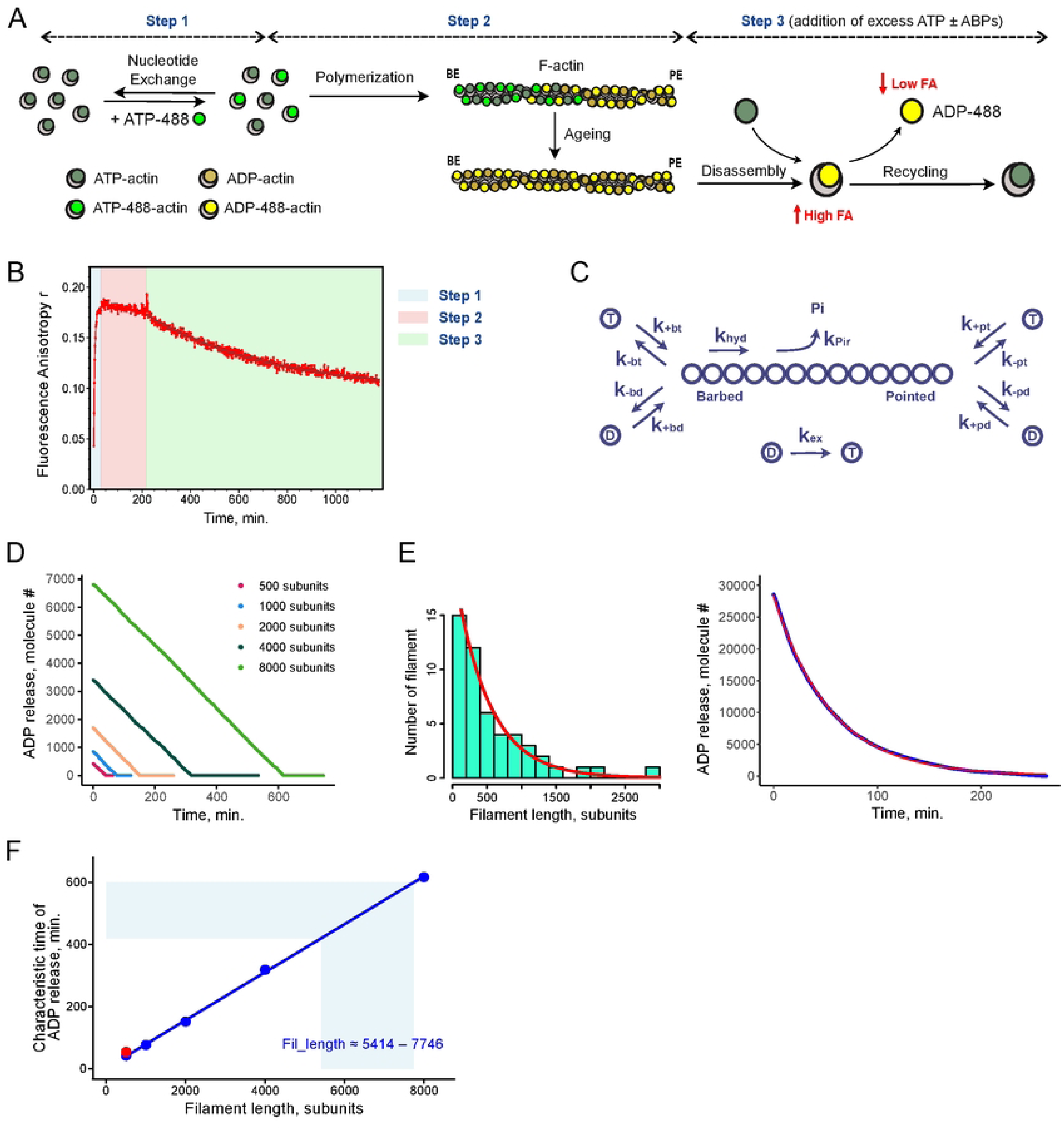
Quantification of actin disassembly and recycling kinetics using a fluorescent nucleotide release assay. **A.** Schematic representation of the assay that measures the rate of fluorescent nucleotide release from F-actin using fluorescence anisotropy. In Step 1, G-actin partially exchanges bound ATP (dark green) for a ATP-488, until steady state is reached. Step 2 corresponds to the polymerization of actin monomers and filament aging when ATP and ATP-488 are progressively hydrolyzed into ADP (dark yellow) and ADP-488 (light yellow). In Step 3, the combined rate of F-actin disassembly and nucleotide exchange on G-actin are measured by fluorescence anisotropy in the presence of excess ATP and in the presence or in the absence of additional ABPs. **B.** Time course of fluorescent nucleotide binding and release from F-actin (2 µM) in the experiment described in A. The anisotropy decay of Step 3 is fitted with a single-exponential function. **C.** Cartoon showing the biochemical reactions that were considered when modelling the actin turnover reaction in silico. **D.** Simulations corresponding to the experiment shown in B. (Step 3), for individual actin filaments of various lengths. Each curve represents the rate of ADP-488 release in solution following depolymerization of ADP-488-bound actin subunits and ADP-488 release from monomers. Rate constants are indicated in Table S1. **E.** Left: Random exponential distribution corresponding to 50 actin filaments with an average length of 500 subunits. Right: Simulation of the kinetics of ADP-488 release in solution for this set of 50 actin filaments (blue) and mono-exponential fit (red). **F.** Characteristic time of fluorescent nucleotide release in solution for simulations presented in D. and E. For single filaments (blue dots), characteristic time is the time interval required until the release of the last fluorescent nucleotide from the filament. For the exponentially distributed filaments (red dot), it is the characteristic time of the mono-exponential fit. Blue line is a linear fit of all points. Light blue area indicates the characteristic time range measured experimentally, and the corresponding filament length distribution.

In the absence of ABPs, corresponding to conditions under which actin filaments undergo treadmilling (Fig. 1B - step 3), the addition of excess ATP leads to a decrease of anisotropy signals over several hours, which can be fitted well as a mono-exponential decay of characteristic time in the 7-10 h range. To identify the processes underlying this decay, we built a stochastic Monte Carlo model of the actin treadmilling reaction, based on published rate constants (Table S1). This model incorporates all relevant chemical reactions, including polymerization and depolymerization at both filament ends, nucleotide hydrolysis, phosphate release and nucleotide exchange on G-actin (Fig. 1C). This last nucleotide exchange reaction on G-actin is considered irreversible from the ADP (or ADP-488) state to ATP due to the presence of excess ATP in solution. For filament populations of uniform length, the simulations predict a constant rate of nucleotide-release until all fluorescent nucleotides have been liberated into solution (Fig. 1D). We interpret this result as a regular replacement of all ADP-488-bound filament subunits by ATP-bound actin monomers during treadmilling. Although individual association and dissociation events occur at both ends, a global net flux of actin monomers is directed towards barbed ends, with an opposite net flux of subunits dissociating from pointed ends. For a filament population of exponential length distribution (Fig. 1E, left), as previously observed in vitro [27], simulations predict an exponential time course of nucleotide release (Fig. 1E, right). Ultimately, these simulations enable us to interpret our experiments as a continuous actin treadmilling reaction occurring within filament populations that follow an exponential length distribution, with an average filament length in the 5400 - 7750 subunit range, corresponding to 16 - 23 µm (Fig. 1F).

### Distinct effects of ABPs on actin disassembly and recycling

We used the same assay to characterize the effect of ADF/cofilin. At concentrations below 1-2 µM, ADF/cofilin promoted up to 12-fold increases in the disassembly and recycling rate of 2 µM F-actin (Fig. 2A). At concentrations above 2 µM, ADF/cofilin had on the contrary an inhibitory effect, which at the highest concentrations culminated in fluorescent nucleotide release rates slower than those observed for spontaneous turnover.

**Figure 2.**
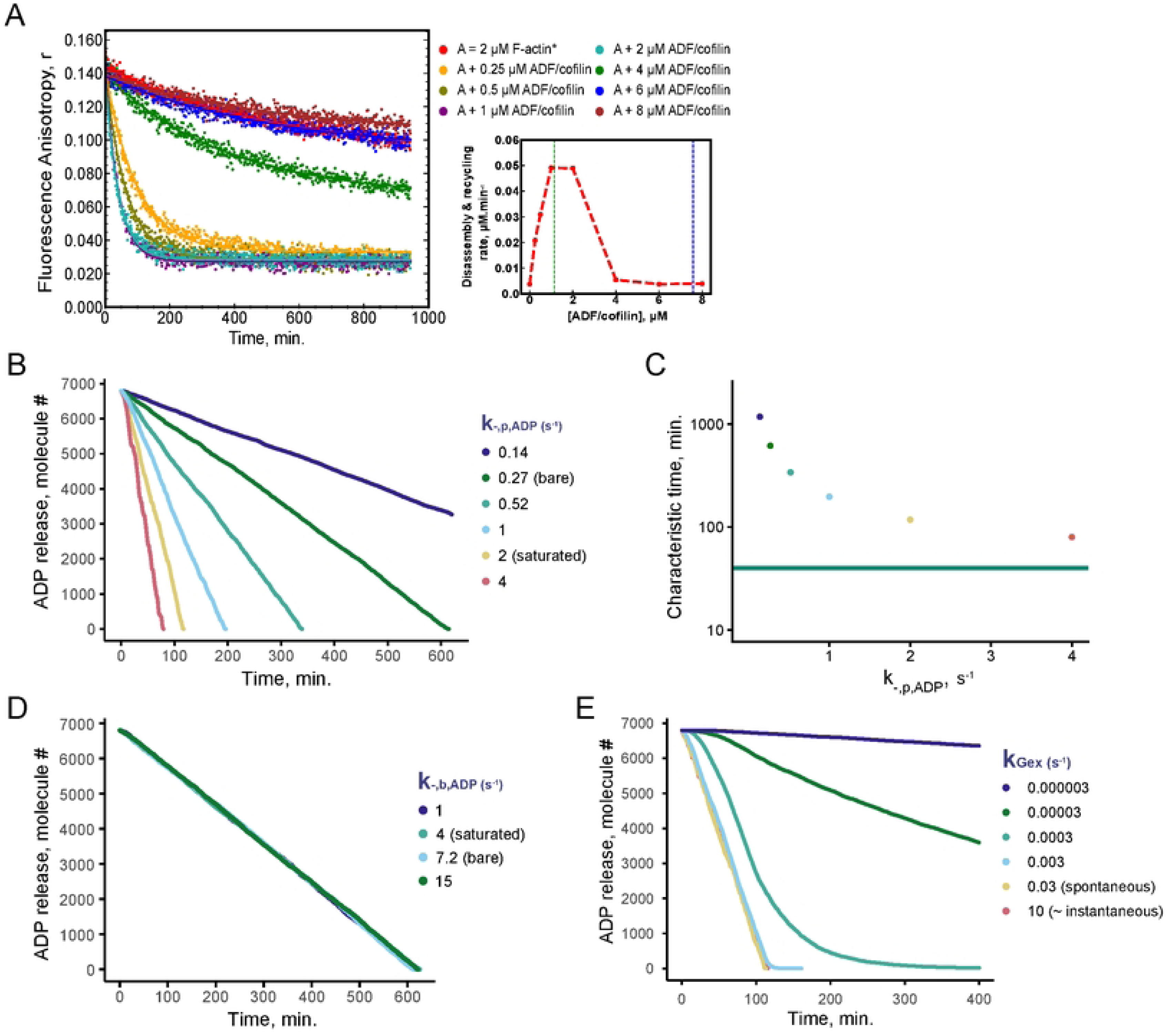
Effect of ADF/cofilin on actin filament disassembly and recycling. **A.** Left: Time course of F-actin (2 µM) disassembly and recycling in the presence of increasing concentration of ADF/cofilin. Right: Quantification of the experiment showing the concentration dependence of ADF/cofilin on disassembly and recycling rates. Blue and green dashed vertical lines indicate the ADF/cofilin concentration and ADF/cofilin concentration equivalent to the ADF/cofilin-to-actin ratio measured in budding yeast [31], respectively. **B.** Simulations of the release time of ADP-488 in solution for 8000 subunit-long actin filaments, when the dissociation constant of ADP subunits from pointed ends 𝑘_―,𝑝,𝐴𝐷𝑃_ varies. Other rate constants are indicated in Table S1. **C.** Quantification of B, indicating with the same color code the characteristic times of ADP-488 release as a function of 𝑘_―,𝑝,𝐴𝐷𝑃_. For comparison, green horizontal lines represents the shortest characteristic time measured experimentally in A. (41 min for 2 µM F-actin + 1 µM ADF/cofilin). **D.** Simulations of the release time of ADP-488 in solution for 8000 subunit-long actin filaments, when the dissociation constant of ADP subunits from actin filament at barbed ends 𝑘_―,𝑏,𝐴𝐷𝑃_varies. Other rate constants are indicated in Table S1. **E.** Simulations of the release time of ADP-488 in solution for 8000 subunit-long actin filaments, when 𝑘_―,𝑝,𝐴𝐷𝑃_ = 𝑘_―,𝑝,𝐴𝐷𝑃,𝑐𝑜𝑓_ = 2 𝑠^―1^ and when the constant rate of nucleotide exchange of ADP for ATP on actin monomers 𝑘_𝐺 𝑒𝑥_ varies. Other rate constants are indicated in Table S1.

How ADF/cofilin’s diverse activities — including fragmentation, subunit depolymerization from filament ends, and nucleotide exchange inhibition — contribute to this process remains unclear. To clarify ADF/cofilin’s effect, we investigated how variations in individual key parameters from earlier simulations impact nucleotide release in solution:

– *Fragmentation activity:* ADF/cofilin fragmentation activity, reflected by changes in filament length in our simulations, clearly provides an efficient means of accelerating the process (Fig. 1D,E).
– *Acceleration of depolymerization:* We varied the dissociation constants of the ADP-bound actin subunits at both ends of the filament over a range of values that covered both spontaneous and ADF/cofilin-induced subunit dissociation [18,28] (Fig. 2B-D). At barbed ends, our simulations indicated that higher dissociation rate constants do not have any significant effect. This is because the frequent association of ATP-bound actin monomers at barbed ends limits any effective dissociation of ADP-bound subunits from these ends (Fig. 2D). In contrast, higher dissociation rate constants at pointed ends lead to faster disassembly and recycling rates (Fig. 2B,C). However, even for the highest dissociation constants published for filaments fully decorated with ADF/cofilin [28], accelerated depolymerization without fragmentation is not sufficient to explain our experimental data (Fig. 2C). Moreover, these simulations overestimate the contribution of depolymerization for two reasons. The first reason is that at these concentrations, ADF/cofilin does not fully decorate F-actin and depolymerization if therefore probably slower. The second reason is that a fraction of ADF/cofilin binds to G-actin, thereby inhibiting nucleotide exchange.
– *Inhibition of nucleotide exchange:* ADF/cofilin strongly inhibits this exchange reaction to levels that are difficult to quantify precisely for actin monomers fully bound to ADF/cofilin. However, based on our previous experiments conducted under similar conditions [26], we estimate that the exchange reaction is at least 100 times slower for G-actin bound to ADF/cofilin. We varied the rate of nucleotide exchange on actin monomers in the simulations, and found that such strong inhibition of nucleotide release from G-actin can prevent the release of fluorescent nucleotides at these experimental time scales (Fig. 2E).

Overall, these simulations indicate that of all ADF/cofilin activities, two of them contribute significantly to actin disassembly (fragmentation and depolymerization at pointed ends), while another one (depolymerization at the barbed ends) contributes negligibly. In contrast, inhibition of nucleotide exchange strongly blocks actin recycling at high concentrations of ADF/cofilin.

These experiments confirm that ADF/cofilin alone drives actin disassembly and monomer recycling too slowly to account for cellular rates [29,30]. They support the involvement of supplementary factors capable of catalyzing all three reactions of fragmentation, depolymerization, and nucleotide exchange at levels that exceed the activity of ADF/cofilin alone. We compared the efficiency of four other proteins in disassembling and recycling of 2 µM F-actin (twinfilin - Fig. 3A, Cyclase Associated Protein (CAP) - Fig. 3B, Aip1 - Fig. 3C and coronin - Fig. 3D). We screened a wide range of concentrations spanning values below and above both the cytoplasmic concentrations (dashed blue lines) and concentrations equivalent to the ABP-to-actin ratio measured in budding yeast (dashed green lines; [31]). Apart from Aip1, all proteins had an effect, albeit with different efficiency. At peak efficiency, ADF/cofilin and twinfilin were the most active proteins, while coronin and CAP were less active. All proteins also showed specific concentration-dependance profiles; coronin, like ADF/cofilin, is particularly effective at low concentrations but has an inhibitory effect at higher concentrations (Fig. 3D - inset); twinfilin’s efficiency increased continuously for all concentrations tested (Fig. 3A - inset); CAP’s efficiency increased and then plateaued (Fig. 3B - inset). Of all the conditions tested, the fastest rate was measured in the presence of 8 µM twinfilin. In this condition, F-actin was disassembled and recycled in 25.2 +/- 1.7 minutes, which is still much slower than what is measured in cells [29,30].

**Figure 3.**
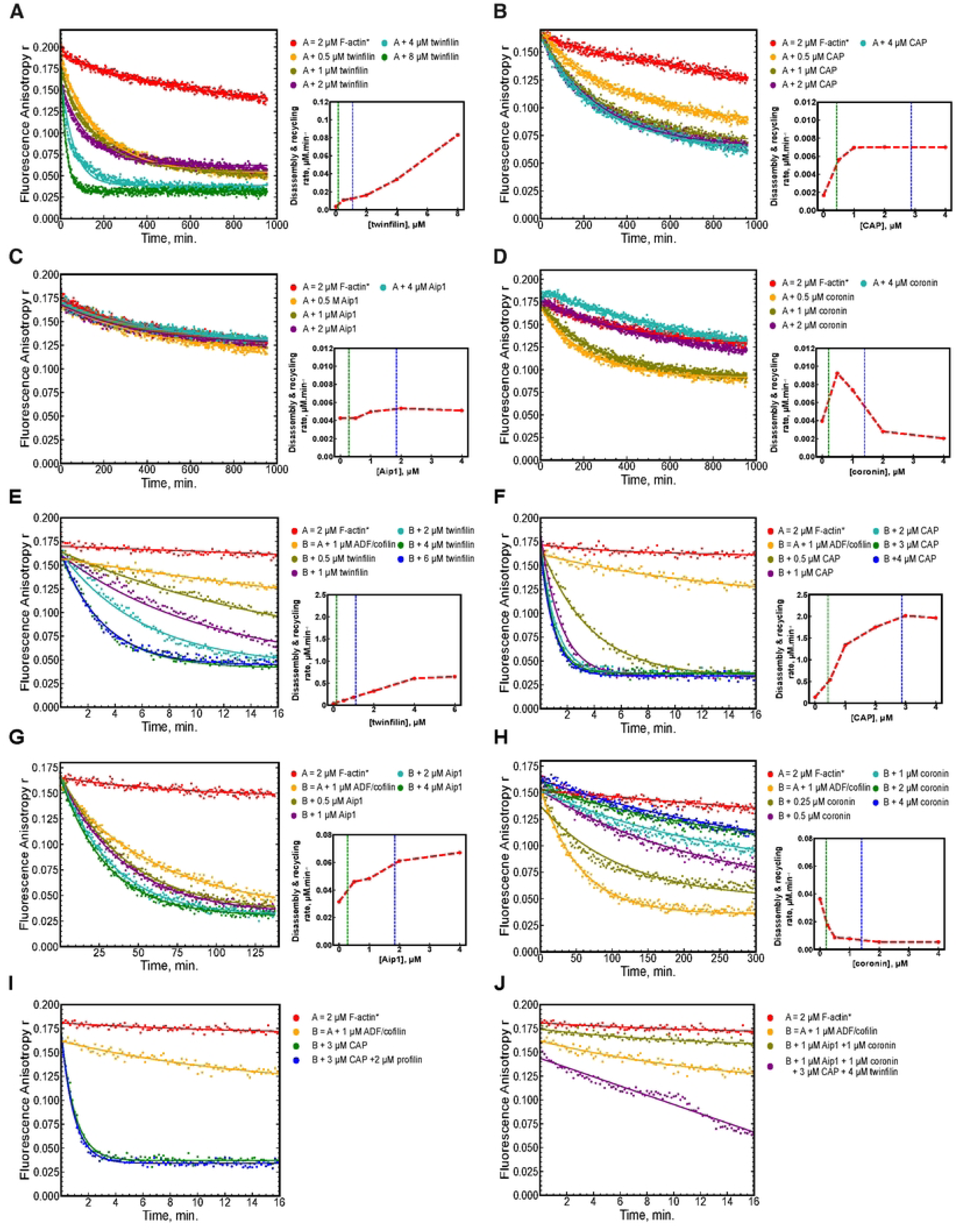
Individual and combinatorial effect of ABPs on F-actin disassembly and recycling. For each panels, left graphs show the time course of F-actin (2 µM) disassembly and recycling measured under the indicated conditions. Right graphs quantify the main panels and show the concentration dependence of the corresponding ABPs on disassembly and recycling rates. Blue and green dashed vertical lines indicate the ABP concentration and ABP concentration equivalent to the ABP-to-actin ratio measured in budding yeast [31], respectively. **A-D.** Effects of increasing concentrations of individual ABPs: twinfilin (in A.), CAP (in B.), Aip1 (in C.) or coronin (in D.). **E-H.** Combinatorial effect of ADF/cofilin (1 µM) with increasing concentrations of twinfilin (in E.), CAP (in F.), Aip1 (in G.) or coronin (in H.). **I.** Effect of 2 µM profilin in the presence of ADF/Cofilin (1 µM) and CAP (3 µM). **J.** Other ABP combinations. The olive-green curve shows the combination of ADF/cofilin, Aip1 and coronin (1 µM each). The purple curve shows the combination of all tested ABPs together: ADF/cofilin (1 µM), Aip1 (1 µM), coronin (1 µM), CAP (3 µM) and twinfilin (4 µM).

### Optimizing conditions for actin disassembly and recycling

We then tested the combinatorial effect between proteins, particularly in the presence of ADF/cofilin, whose activity is enhanced by multiple factors. With 2 µM F-actin, we found that the activity of 1 µM of ADF/cofilin was enhanced by either CAP, twinfilin or Aip1 (Fig. 3E-G). On the contrary, coronin had an inhibitory effect (Fig. 3H). The fastest rates were measured in the presence of 1 µM ADF/cofilin and 6 µM twinfilin (216 +/- 39 seconds) or 1 µM ADF/cofilin and 3 µM CAP (62.7 +/- 5.5 seconds), which correspond more closely to the values expected in non-muscle cells. The addition of profilin to accelerate nucleotide exchange on actin monomers further did not lead to significant improvement (Fig. 3I). The same was true for the more complex combinations we tested (ADF/cofilin + Aip1 + coronin or ADF/cofilin + Aip1 + coronin + twinfilin + CAP) (Fig. 3J). Therefore, combinations of ADF/cofilin and CAP or twinfilin seem optimal and sufficient for reconstituting actin dynamics in vitro at physiological rates.

### Effect of F-actin concentration on rates of actin disassembly and recycling

The activities of actin-regulating proteins have been studied extensively under conditions where actin concentrations are low (on the order of micromolar). The influence of actin concentration on their activity remains largely unknown, particularly at the much higher actin levels found in eukaryotic cells (typically one to two orders of magnitude greater [31–33]). To assess the effect of concentration, we increased here the F-actin concentration from 2 to 10 µM and examined both the activity and concentration-dependence of ADF/cofilin (Fig. 4A,B). Surprisingly, ADF/cofilin was less efficient and its activity less concentration-dependent at this higher F-actin concentration, with no obvious optimal range for efficient activity of ADF/cofilin. Figure 4B, which plots the amount of polymer disassembled per unit of time, reveals that this rate is not entirely conserved between the experiments conducted at 2 and 10 µM, whether the comparison is made at equal ADF/cofilin concentrations, at equal actin:ADF/cofilin ratios, or at peak efficiency. We suspected first that the fragmentation activity of ADF/cofilin, which depends greatly on the flexibility and buckling of actin filaments, may decrease at high F-actin concentrations because of lower degree of filament freedom. However, experiments performed with 2 µM F-actin in the presence of methylcellulose, an agent which increases the viscosity of the medium and promotes the formation of actin bundles, did not indicate any reduction of ADF/cofilin’s activity and do not support this hypothesis (Fig. 4C).

**Figure 4.**
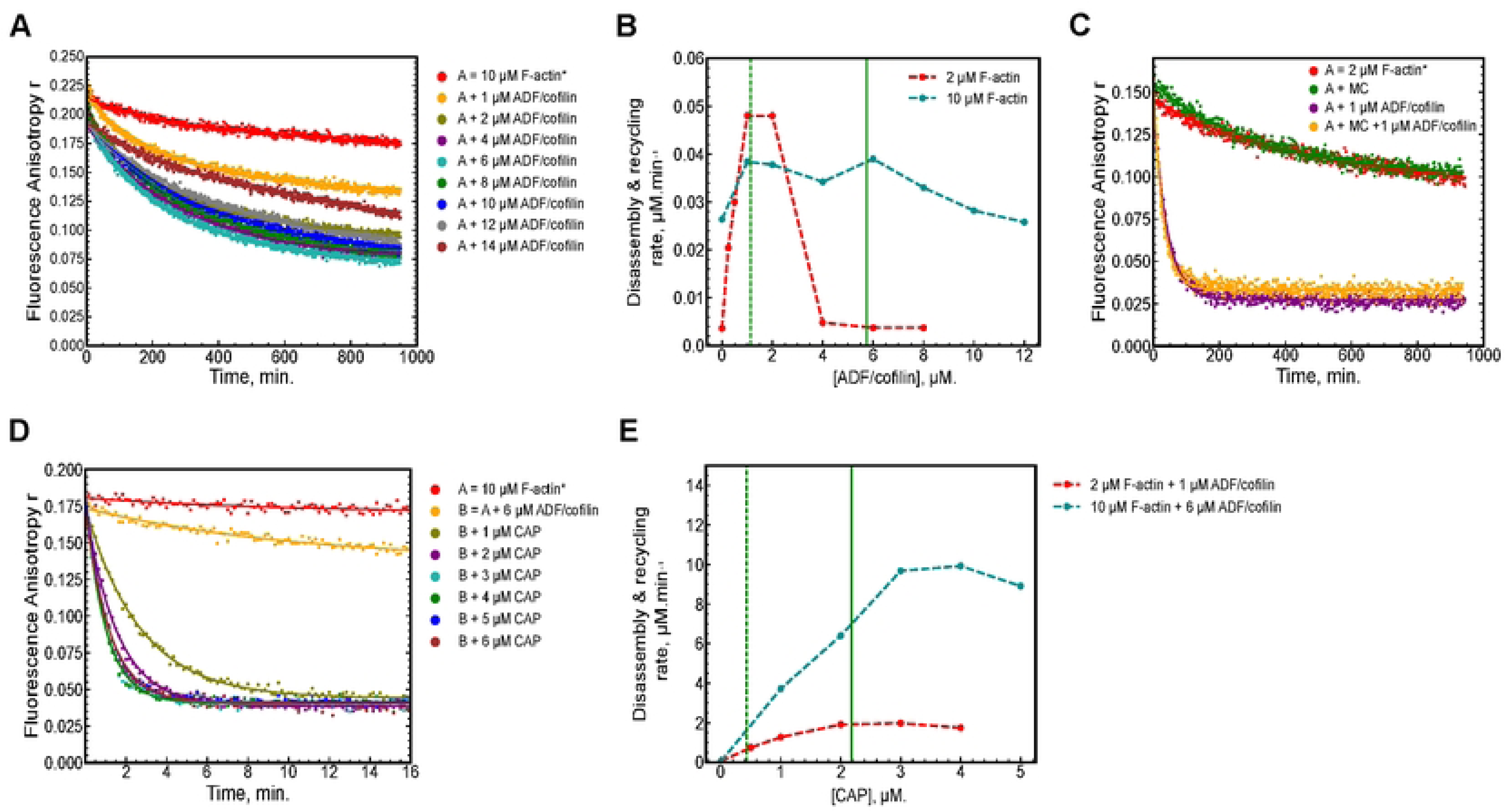
Effect of F-actin concentration on rates of actin disassembly and recycling. **A.** Time course of F-actin (10 µM) disassembly and recycling in the presence of increasing concentration of ADF/cofilin. **B.** Quantification of Fig. 2A and 4A showing the concentration dependence of ADF/cofilin on the disassembly and recycling rates of 2 µM (in red) or 10 µM (in cyan) of F-actin. Green vertical lines indicate the ADF/cofilin-to-actin ratio measured in budding yeast [31] for 2 µM F-actin (dashed line) and for 10 µM F-actin (solid line). **C.** Time course of F-actin (2 µM) disassembly and recycling in 0.3% methylcellulose (MC), in absence or presence of ADF/Cofilin (1 µM). **D.** Time course of F-actin (10 µM) disassembly and recycling in the presence of 6 µM ADF/cofilin and increasing concentration of CAP. **E.** Quantification of Fig. 2F and 4D showing the concentration dependence of CAP on the disassembly and recycling rates of 2 µM F-actin and 1 µM ADF/cofilin (in red) or 10 µM F-actin and 6 µM ADF/cofilin (in cyan). Green vertical lines indicate the CAP-to-actin ratio measured in budding yeast [31] for 2 µM F-actin (dashed line) and 10 µM F-actin (solid line).

We next asked whether specific properties of a protein such as CAP could bypass this limitation. Notably, actin filament disassembly and recycling were markedly more efficient when 10 µM F-actin was incubated with ADF/cofilin and CAP, displaying a clear dependence on CAP concentration. Under optimal conditions, the total amount of polymer disassembled per unit time was roughly five times higher than that observed with 2 µM F-actin, indicating that the reaction rate now scales with the amount of F-actin (Fig. 3E,F). These findings demonstrate that, under these optimal conditions, ADF/cofilin and CAP can drive actin disassembly and recycling efficiently, regardless of the initial actin concentration.

### Measurement of actin turnover rates and energy consumption using an NADH-coupled assay

We next asked whether the identified protein set was sufficient to support rapid actin turnover in vitro. Achieving fast actin dynamics requires not only efficient F-actin disassembly and monomer recycling but also that the released actin monomers are capable of re-polymerizing rapidly. In each turnover cycle, every actin subunit hydrolyzes a single ATP molecule, which is then released as ADP. Thus, the rate of ADP production serves as a dual metric, quantifying both the efficiency of actin turnover and the energy consumption of the system.

The amount of ADP released can be measured by enzymatic methods involving the coupling of an ATP regenerator system, whose by-product is pyruvate, and a pyruvate-to-lactate conversion system that oxidizes one NADH molecule into NAD^+^. Because NADH is auto-fluorescent and NAD^+^ is not, the rate of actin turnover in this system can be measured by quantifying the loss of NADH fluorescence (Fig. 5A, B) [34]. We used this system to quantify the turnover rate of actin (10 µM) at steady-state, and found that ATP consumption increases with the addition of ADF/cofilin and CAP (Fig. 5C). In the presence of ADF/cofilin and CAP, the slope difference between the curve (in red) and the control condition without protein (in blue) allows the ATP consumption of the system to be calculated accurately, thus providing a reliable value for the actin turnover rate (Fig. 5D). We compared this rate to the rate of disassembly and recycling measured in anisotropy experiments under similar protein concentration and buffer conditions (Fig. 4D,E) and found comparable values (Fig. 5D). This result indicates the equality of both rates under conditions of dynamic turnover and validates the complementarity of the two methods.

**Figure 5.**
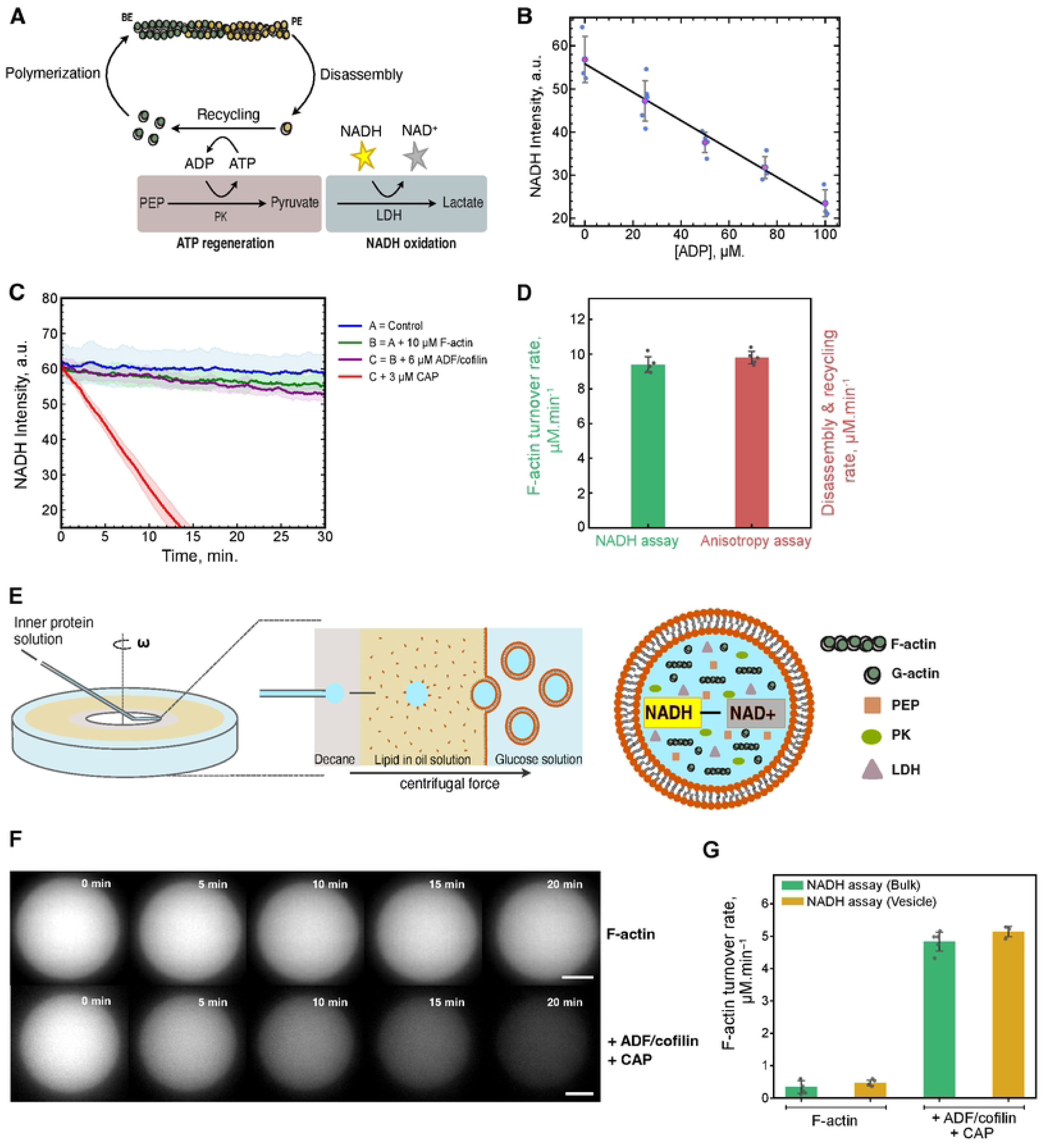
Actin turnover rates and energy consumption measured using an NADH-coupled enzymatic assay in bulk and GUVs. **A.** Schematic diagram of the NADH-coupled enzymatic assay, in which actin turnover rates and energy consumption are quantified by measuring the amount of ADP produced by the system. **B.** Calibration curve relating NADH (200 µM) fluorescence intensity as a function of ADP concentration, in the presence of 0.2 mM PEP, 21 unit.ml^-1^ PK and 31 unit.ml^-1^ LDH (n = 4). Blue dots indicate individual measurements, red dots mean values with standard deviations. Black line is a linear fit of the data. **C.** Time course of NADH fluorescence intensity for a control experiment without F-actin and ABPs (blue line, n = 6), with 10 µM F-actin (green line, n=6), with 10 µM F-actin and 6 µM ADF/cofilin (purple line, n=4), or with 10 µM F-actin, 6 µM ADF/cofilin and 3 µM CAP (red line, n=5). In this experiment, 1 mM ATP was added to match the buffer conditions of the corresponding anisotropy experiment (from Fig. 4D). For each condition, solid lines represent the mean NADH intensity and the shaded regions indicate the standard deviation at each time point. **D.** Comparison of the actin turnover rate calculated from the NADH-coupled assay (from Fig. 5C), with the disassembly and recycling rate calculated from the corresponding fluorescence anisotropy assay (from Fig. 4D), in presence of ADF/cofilin and CAP (n=5 for both assays). **E.** Schematics of the cDICE method used to encapsulate GUVs with F-actin, ABPs and the NADH-coupled assay components (inner protein solution). Left part illustrates how droplets of these mixtures are injected into the cDICE chamber, which is filled with three different fluids. The central section is a cross-sectional view showing the droplet passing through several interfaces until a GUV is formed. The right-hand side shows the components that are finally encapsulated inside the GUV. **F.** Snapshots of NADH fluorescence from GUVs measured over 20 minutes. Top panel shows a GUV encapsulated with 10 µM F-actin (scale bar = 10 µm) and bottom panel shows a GUV encapsulated with 10 µM F-actin, 6 µM ADF/cofilin and 3 µM CAP along with NADH-coupled assay components (scale bar = 15 µm). **G.** Comparison of actin turnover rates measured in bulk solutions and in GUVs. n=6 (F-actin in bulk); n=5 (F-actin + ADF/cofilin + CAP in bulk); n=5 (F-actin in GUVs); n= 3 (F-actin + ADF/cofilin + CAP in GUVs). Contrary to experiments of panel C, no additional ATP was provided here.

Finally, we wanted to determine whether the rapid actin turnover kinetics measured in bulk are maintained in cell-sized experimental systems. Indeed, when the size of a biochemical system changes, depletion effects or changes in protein stoichiometry can alter the kinetics of certain biochemical reactions [25]. We used cDICE encapsulation method to internalize the same reaction mixtures inside giant unilamellar vesicles (GUVs), and followed rates of NADH fluorescence decrease by fluorescence microscopy [35]. Our measurements show that reaction rates in bulk and within GUVs are consistently identical, demonstrating that rapid actin turnover can be faithfully reconstituted at high actin concentrations inside cell-sized GUVs with this protein combination, and that NADH-coupled assays enable precise turnover rate quantification by fluorescence microscopy.

## Discussion

This work provides the first side-by-side comparison of the efficiency of ABPs implicated in actin disassembly and turnover. We have established new high-throughput approaches that enable rapid investigation of protein activity across physiologically relevant concentration ranges and ratios. Building upon a large number of results previously published in the literature, this study provides comparative data, mechanistic details and optimized experimental conditions of actin turnover for future actin-based reconstitutions in cell-sized systems.

In the case of ADF/cofilin, its strong inhibition of nucleotide exchange at high concentration is problematic and requires the involvement of additional enzymes such as profilin or CAP to catalyze nucleotide exchange. We also found that both accelerated depolymerization at pointed ends and fragmentation contribute significantly to the disassembly of F-actin by ADF/cofilin, but are not sufficient on their own. The intervention of additional factors, such as CAP or twinfilin, is essential to achieve disassembly at rates such as those observed in cells. These proteins act by increasing greatly the rate of actin depolymerization, or by more efficient mechanisms such as bursting. Our study also shows that the efficiency of the process varies greatly depending on the protein families involved and their concentration-dependence. For certain proteins such as Aip1 and coronin, the activities we measure may appear to be lower than those observed in previous studies. However, a direct comparison is difficult because most published studies analyze disassembly or recycling processes independently, whereas our method analyzes the combination of both. Other experimental differences, such as variability between protein homologues, may also explain differences. In this study, we intentionally used rabbit muscle actin and *S. cerevisiae* ABPs, as these are the most commonly used proteins in reconstituted assays, and for which the corresponding rate constants and affinities have been measured. However, it would be important to investigate in the future how homologous proteins differ.

Another notable point highlighted by this study is the non-conservation of ABP efficiency under various concentration of F-actin. Our findings reveal a strong and non-intuitive dependence on this concentration factor, which often differs substantially between in vitro experiments and the cellular conditions. At the two F-actin concentrations tested here, ADF/cofilin alone tends to disassemble and recycle at a relatively constant polymer quantity per unit of time, while a set of proteins such as ADF/cofilin + CAP can increase disassembly and recycling rates according to the initial polymer quantity. One reason for this effect probably comes from the mechanism of ADF/cofilin itself, whose binding to F-actin is highly cooperative and whose disassembly efficiency depends on the degree of protein binding. Thus, the formation of ADF/cofilin domains of optimal length along actin filaments may depend more on the concentration of ADF/cofilin itself than on the concentration of F-actin. If true, this suggests complex stoichiometry effects between F-actin and ADF/cofilin, where the kinetics of a reaction at X µM F-actin + Y µM ADF/cofilin do not allow for a trivial prediction of the kinetics of the reaction at (n·X) µM F-actin + (n·Y) µM ADF/cofilin. More advanced modeling of these reactions will be necessary to understand these effects.

We also demonstrated that NADH-coupled assays could be implemented to quantify the energy consumption of F-actin solutions and the influence of ABPs. Because each actin molecule that polymerizes and integrates into filaments consumes one ATP molecule, these tests also reflect the rate of actin turnover. Importantly, values of disassembly/recycling rates measured by fluorescence anisotropy are equal to those obtained from NADH-coupled turnover rate assays, thus validating both approaches. In the best conditions (10 µM F-actin, 6 µM ADF/cofilin and 4 µM CAP), we measure actin turnover rates of about 10 µM.min^-1^, indicating that each actin molecule goes through one polymerization cycle and consumes 1 ATP per minute, which reflects well cellular observations and seems to indicate that optimal experimental conditions may have been reached here.

## Acknowledgments

This project has received funding from the Fondation pour la Recherche Médicale to AM, grant « Équipe FRM EQU202103012764 ».

## Methods

### Protein purification

Actin was purified from rabbit muscle acetone powder as described in [36], and stored in G-buffer (5 mM Tris pH 8; 0.1 mM CaCl_2_; 0.2 mM ATP; 0.5 mM DTT and 0.02% NaN_3_).

ADF/cofilin (Uniprot # Q03048) was expressed in a Rosetta2 (DE3) bacterial strain and purified as described in [37]. Twinfillin (Uniprot # P53250), Coronin (Uniprot # Q06440) and Aip1 (Uniprot # P46680) were cloned, expressed and purified as described for Aip1 in [14].

CAP (Uniprot # P17555) was cloned as a 6xHis-SUMO-TEV N-terminal construct, then expressed and purified in an ArcticExpress (DE3) bacterial strain. Bacteria were grown in Terrific buffer at 37°C up to OD = 0.4, then at 13°C up to OD = 0.47, before induction with 0.25 mM IPTG for 46 h. Cells were harvested by centrifugation and flash frozen in liquid nitrogen. Pellets were resuspended in lysis buffer (50 mM Tris, pH 7.5; 250 mM NaCl; 10 mM MgCl_2_) with 0.5 mg/ml lysozyme (Euromedex #5933-D), 20ug/mL DNAseI (Roche #11284932001) and cOmplete EDTA-free protease inhibitors (Roche). Cells were sonicated and centrifugated for 30 min at 40,000 rpm. Cleared supernatants were incubated for 2h at 4°C with 4 ml of Ni beads (Ni Sepharose6 FastFlow (Cytiva #17531802)). Beads were washed at low Imidazole (50 mM Tris, pH 7.5; 250 mM NaCl; 25 mM Imidazole) and eluted at high Imidazole (50 mM Tris, pH 7.5; 250 mM NaCl; 250 mM Imidazole). TEV protease was added to cleave the purification tag, cleaved protein was dialyzed overnight in (50 mM Tris, pH 7.5; 250 mM NaCl; 0.5 mM TCEP) and purification tags were eliminated with 4 ml of Ni beads. Proteins were finally concentrated down to 600 µl, aliquoted, flash frozen and stored at -80°C. Their concentration was measured from their optical density at 280 nm and their predictive molar extinction coefficient estimated from Expasy-ProtParam.

### Fluorescence anisotropy assays

The fluorescent nucleotide analogue used in this study is N^6^-(6-Amino)hexyl-ATP-ATTO-488 (referred to as ATP-488; Jena Bioscience ref. NU-805-488). Stock solutions were diluted to 10 μM in 20 mM Hepes pH 7.5, aliquoted and stored at -80°C to avoid repeated freeze/thaw cycles. Experiments on actin disassembly and recycling by fluorescence anisotropy assays were performed in three sequential steps (steps 1 to 3), all conducted at room temperature. During step 1 (partial exchange of ATP for ATP-488), G-actin was incubated with ATP-488 for 30 min in a nucleotide-free G-buffer (NFG buffer; 5 mM Tris pH 8; 0.1 mM CaCl_2_; 0.5 mM DTT and 0.02% NaN_3_), supplemented with 10x ME (1x ME contains 1 mM MgCl_2_, 1 mM EGTA). During step 2 (actin polymerization), actin monomers were polymerized for 2 hours into filaments upon addition of 50 mM KCl. During step 3 (release of fluorescent nucleotide from actin), an excess of ATP (1 mM) was added to the reaction mixture. As the addition of ATP initially caused a slight decrease in the anisotropy signals during the first few minutes, due to the release of fluorescent nucleotides from remaining actin monomers in solution at critical concentrations, ABPs were added to the reaction mixture 10 minutes after the ATP. Fluorescence anisotropy kinetics were recorded on a Safas Xenius XC spectrofluorometer, using a multi-cuvette mode. ATP-488 was excited at 504 nm and the emitted light was collected at 521 nm, with both bandwidths set to 10 nm.

#### Data Analysis

All fluorescence anisotropy data were plotted and fitted with a single-exponential function using Python. The disassembly and recycling rates were calculated and plotted as the product [𝐹 ― 𝑎𝑐𝑡𝑖𝑛] · 𝑟_𝑜𝑓𝑓,𝐴𝐷𝑃―488_ (in μM·s^-1^), where 𝑟_𝑜𝑓𝑓,𝐴𝐷𝑃―488_ is the rate of fluorescent nucleotide dissociation obtained from curve fitting, and [𝐹 ― 𝑎𝑐𝑡𝑖𝑛] is the total actin concentration used in the assay.

### Protein encapsulation in GUVs by cDICE technique

#### Lipid-in-oil solution (LOS) preparation

Stock solutions of L-α-phosphatidylcholine (EggPC; Sigma-Aldrich P3556), 1,2-Distearoyl-sn-glycero-3-phosphoethanolamine-N-[methoxy(polyethyleneglycol)-2000] (18:0 PEG2000 PE; Avanti Polar 880120C) and 1,2-distearoyl-sn-glycero-3-phosphoethanolamine-N-[biotinyl(polyethylene glycol)-2000] (DSPE-PEG(2000)-Biotin; Avanti; 880129C) were stored in chloroform at -20 °C. For each experiment, 5 ml of LOS was prepared by mixing lipids in 300 μl decane, followed by dropwise addition of 4.7 ml of a silicone oil (Carl Roth 4020 – 50 cSt) /mineral oil (Sigma-Aldrich M3516) mixture (85:15; v/v) under gentle vortexing. The final concentration of lipids in LOS was 0.5 mM, with standard lipid composition of 95% of EggPC and 5% of PEG2000-PE. For biotinylated lipid mixtures 1% DSPE-PEG(2000)-Biotin was added by reducing EggPC fraction to 94%.

#### Encapsulated Aqueous Solution (EAS) preparation

The EAS contains 300 mM of sucrose, NADH, PEP, PK/LDH, F-actin and ABPs in G-buffer supplemented with KME. F-actin and ABPs were added to the EAS only after filling the cDICE chamber and immediately before loading into the capillary

#### Dispersing aqueous solution (DAS) preparation

The DAS contains glucose dissolved in water. The concentration of glucose in the DAS was adjusted to reach an osmolarity 5-15 m Osm higher than the EAS.

#### Capillaries preparation

Glass capillaries with inner diameter 0.58 mm and outer diameter 1 mm (WPI; B100-58-15) were pulled using a pipette puller (model P1000 Sutter Instrument), and pipette tips were cut to a diameter of 15-20 μm with a microforge. Capillaries were bent using a flame to facilitate insertion into the horizontal chamber, and tips were silanized in 2 % trichloro(octadecyl)silane in toluene. Capillaries were stored under dry conditions.

#### GUV production

GUVs were prepared at room temperature using a home-built cDICE setup. A custom-fabricated plexiglass chamber with a 3D-printed resin lid was mounted horizontally on a motor and rotated at 1800 rpm. The chamber was sequentially filled with 1-1.2 ml DAS, 3-4 ml LOS and 0.5-0.7 CP (Decane). Capillaries were loaded with EAS, connected to a pressure device (150-200 mBar), and the tip was inserted into the decane layer. The chamber rotation was stopped after 5-20 minutes of droplet injection. The glucose phase containing the vesicles was collected, and GUVs sedimented by gravity in an Eppendorf tube for 2-3 minutes. *GUV immobilization and imaging:* Glass coverslips were sonicated in ethanol for 15 min, rinsed with Milli-Q water, and dried prior to use. They were incubated with a PBS solution containing BSA (Sigma-Aldrich A9418; 0.7 mg.ml^-1^) and BSA-biotin (Sigma-Aldrich; A8549; 0.3 mg.ml^-1^) for 20 minutes at room temperature. After two washes in PBS, they were incubated with 0.5 mg.ml^-1^ streptavidin for 20 minutes. The chamber was rinsed with DAS. 5 mM of KCl was added to the GUVs to reduce short-range electrostatic repulsion and facilitate the streptavidin-binding interaction.

### GUVs imaging and analysis

Images of the GUVs were acquired using a Zeiss Axio Observer Z1 inverted wide-field microscope, which was equipped with a 60x/1.4 NA oil objective and a Hamamatsu ORCA-Flash 4.0 LT camera. NADH fluorescence within the GUVs was recorded using Zen 2.3 (Blue Edition) software. For NADH excitation and emission, a DAPI filter set was used. Images were analyzed using ImageJ software. The fluorescence intensity of each vesicle was subtracted by the background intensity.

### NADH-coupled enzymatic assays

#### Experiments in bulk

Nicotinamide adenine dinucleotide (NADH; 0.2 mM; Targetmol T5283), phosphoenolpyruvate (PEP; 0.2 mM; Sigma-Aldrich ref. P7127) and a pyruvate kinase/lactate dehydrogenase mix (PK/LDH; 21/31 unit.ml^-1^; Sigma-Aldrich P0294) were mixed with 10 µM F-actin and indicated ABPs in NFG buffer supplemented with 10x KME (1x KME contains 50 mM KCl; 1 mM MgCl_2_; 1 mM EGTA) and indicated concentration of ATP. The decrease in NADH fluorescence was recorded immediately using a Safas Xenius XC spectrofluorometer. Excitation was set to 329 nm, and emission light was collected at 451 nm. The excitation and emission bandwidths were set to 10 and 15 nm, respectively.

#### Experiments in GUVs

The same NADH assay components were encapsulated into GUVs, except that the NADH concentration was increased to 0.4 mM to maintain a strong fluorescence signal inside the GUVs at the start of imaging. It was confirmed using bulk assay that 0.2 mM of PEP and 21/31 unit.ml^-1^ PK/LDH enzymes are sufficient to support the oxidation of 0.4 mM NADH for the duration of the imaging.

### Actin turnover rate calculation in bulk and inside GUVs

ADP production was calibrated by measuring the fluorescence intensity of NADH in the presence of known concentrations of ADP in both bulk and vesicles. The decrease in fluorescence intensity was corrected for photobleaching by subtracting the percentage loss of NADH intensity measured under identical experimental conditions without F-actin. The rate of change of ADP concentration was used to determine the actin turnover rate.

### Rejection-free Monte Carlo method to simulate the turnover of actin

Stochastic simulations use a Monte Carlo method [38], implemented with a rejection-free algorithm similar to the Gillespie algorithm [39]. In these simulations, all polymerization/depolymerization, random nucleotide hydrolysis and random phosphate release are treated as probabilities for these events to occur. The biochemical parameters used to run these models are listed in Table S1.

The algorithm works by selecting the next molecular event at each step, based on its probability of occurrence. The model runs as follows at each step:

1. The transition rates 𝑟_𝑖_for each reaction at both filament ends are calculated. For the association of actin monomers, they are equal to 𝑘_+,𝐴𝑇𝑃_ · [𝐺 ― 𝐴𝑇𝑃] and 𝑘_+,𝐴𝐷𝑃_ · [𝐺 ― 𝐴𝐷𝑃]. For the dissociation of actin subunits, they are equal to 𝑘_―,𝐴𝑇𝑃_ and 𝑘_―,𝐴𝐷𝑃_.
2. The sum of the transition rates 𝑟_𝑂_ = ∑_𝑖_ 𝑟_𝑖_ is calculated.
3. One of the possible events is selected randomly, based on its probability 𝑟_𝑖_/𝑟_0_ to occur
4. The state of the filament, pool of monomers and ADF/cofilin are updated.
5. We pick a uniform number u ∈ (0,1) and update the time 𝑡 = 𝑡 + Δ𝑡 (initial time is t = 0), where 𝑡 = ― log ^𝑢^/_𝑟_0__
6. We proceed to random hydrolysis of ATP-bound subunits, random phosphate release of ADP-Pi-bound subunits and random nucleotide exchange on ADP-bound actin monomers based on the probability of these events occurring during Δ𝑡.
7. This process is repeated until the system reaches steady-state

Simulations track the fate of ADP nucleotides pre-bound to F-actin until their disassembly and release from actin monomers.

## Supporting information caption

**Table S1:** Biochemical parameters used to simulate the turnover of actin

## References

1. Goode BL, Eskin J, Shekhar S. Mechanisms of actin disassembly and turnover. Journal of Cell Biology. 2023;222. doi:10.1083/jcb.202309021

2. Lappalainen P, Kotila T, Jégou A, Romet-Lemonne G. Biochemical and mechanical regulation of actin dynamics. Nat Rev Mol Cell Biol. 2022;23: 836–852. doi:10.1038/s41580-022-00508-4

3. Skau CT, Waterman CM. Specification of Architecture and Function of Actin Structures by Actin Nucleation Factors. Annu Rev Biophys. 2015;44: 285–310. doi:10.1146/annurev-biophys-060414-034308

4. Pollard TD. Actin and Actin-Binding Proteins. Cold Spring Harb Perspect Biol. 2016;8: a018226. doi:10.1101/cshperspect.a018226

5. Colin A, Kotila T, Guérin C, Orhant-Prioux M, Vianay B, Mogilner A, et al. Recycling of the actin monomer pool limits the lifetime of network turnover. EMBO J. 2023;42. doi:10.15252/embj.2022112717

6. Gandhi M, Smith BA, Bovellan M, Paavilainen V, Daugherty-Clarke K, Gelles J, et al. GMF Is a Cofilin Homolog that Binds Arp2/3 Complex to Stimulate Filament Debranching and Inhibit Actin Nucleation. Current Biology. 2010;20: 861–867. doi:10.1016/j.cub.2010.03.026

7. Koundinya N, Aguilar RM, Wetzel K, Tomasso MR, Nagarajan P, McGuirk ER, et al. Two ligands of Arp2/3 complex, yeast coronin and GMF, interact and synergize in pruning branched actin networks. Journal of Biological Chemistry. 2025;301. doi:10.1016/j.jbc.2025.108191

8. Michelot A, Berro J, Guérin C, Boujemaa-Paterski R, Staiger CJ, Martiel JL, et al. Actin-Filament Stochastic Dynamics Mediated by ADF/Cofilin. Current Biology. 2007;17: 825–833. doi:10.1016/j.cub.2007.04.037

9. Andrianantoandro E, Pollard TD. Mechanism of Actin Filament Turnover by Severing and Nucleation at Different Concentrations of ADF/Cofilin. Mol Cell. 2006;24: 13–23. doi:10.1016/j.molcel.2006.08.006

10. De La Cruz EM, Sept D. The kinetics of cooperative cofilin binding reveals two states of the cofilin-actin filament. Biophys J. 2010;98: 1893–1901. doi:10.1016/j.bpj.2010.01.023

11. Jégou A, Niedermayer T, Orbán J, Didry D, Lipowsky R, Carlier MF, et al. Individual actin filaments in a microfluidic flow reveal the mechanism of ATP hydrolysis and give insight into the properties of profilin. PLoS Biol. 2011;9. doi:10.1371/journal.pbio.1001161

12. Carlier MF, Pantaloni D. Direct evidence for ADP-inorganic phosphate-F-actin as the major intermediate in ATP-actin polymerization. Rate of dissociation of inorganic phosphate from actin filaments. Biochemistry. 1986;25: 7789–7792. doi:10.1021/bi00372a001

13. Suarez C, Roland J, Boujemaa-Paterski R, Kang H, McCullough BR, Reymann AC, et al. Cofilin tunes the nucleotide state of actin filaments and severs at bare and decorated segment boundaries. Current Biology. 2011;21: 862–868. doi:10.1016/j.cub.2011.03.064

14. Gressin L, Guillotin A, Guérin C, Blanchoin L, Michelot A. Architecture dependence of actin filament network disassembly. Current Biology. 2015;25: 1437–1447. doi:10.1016/j.cub.2015.04.011

15. Hao YK, Charras GT, Mitchison TJ, Brieher WM. Actin disassembly by cofilin, coronin, and Aip1 occurs in bursts and is inhibited by barbed-end cappers. Journal of Cell Biology. 2008;182: 341–353. doi:10.1083/jcb.200801027

16. Oosterheert W, Boiero Sanders M, Hofnagel O, Bieling P, Raunser S. Choreography of rapid actin filament disassembly by coronin, cofilin, and AIP1. Cell. 2025;188: 6845–6860.e27. doi:10.1016/j.cell.2025.09.016

17. Oosterheert W, Boiero Sanders M, Bieling P, Raunser S. Structural insights into actin filament turnover. Trends in Cell Biology. Elsevier Ltd; 2025. pp. 893–906. doi:10.1016/j.tcb.2024.12.009

18. Shekhar S, Chung J, Kondev J, Gelles J, Goode BL. Synergy between Cyclase-associated protein and Cofilin accelerates actin filament depolymerization by two orders of magnitude. Nat Commun. 2019;10. doi:10.1038/s41467-019-13268-1

19. Kotila T, Wioland H, Enkavi G, Kogan K, Vattulainen I, Jégou A, et al. Mechanism of synergistic actin filament pointed end depolymerization by cyclase-associated protein and cofilin. Nat Commun. 2019;10. doi:10.1038/s41467-019-13213-2

20. Jansen S, Collins A, Chin SM, Ydenberg CA, Gelles J, Goode BL. Single-molecule imaging of a three-component ordered actin disassembly mechanism. Nat Commun. 2015;6. doi:10.1038/ncomms8202

21. Nadkarni A V., Brieher WM. Aip1 destabilizes cofilin-saturated actin filaments by severing and accelerating monomer dissociation from ends. Current Biology. 2014;24: 2749–2757. doi:10.1016/j.cub.2014.09.048

22. Shekhar S, Hoeprich GJ, Gelles J, Goode BL. Twinfilin bypasses assembly conditions and actin filament aging to drive barbed end depolymerization. Journal of Cell Biology. 2021;220. doi:10.1083/jcb.202006022

23. Johnston AB, Collins A, Goode BL. High-speed depolymerization at actin filament ends jointly catalysed by Twinfilin and Srv2/CAP. Nat Cell Biol. 2015;17: 1504–1511. doi:10.1038/ncb3252

24. Wioland H, Egou AJ, Romet-Lemonne G. Celebrating 20 years of live single-actin-filament studies with five golden rules. doi:10.1073/pnas.2109506119/-/DCSupplemental

25. Kandiyoth FB, Michelot A. Reconstitution of actin-based cellular processes: Why encapsulation changes the rules. Eur J Cell Biol. 2023;102: 151368. doi:10.1016/j.ejcb.2023.151368

26. Colombo J, Antkowiak A, Kogan K, Kotila T, Elliott J, Guillotin A, et al. A functional family of fluorescent nucleotide analogues to investigate actin dynamics and energetics. Nat Commun. 2021;12. doi:10.1038/s41467-020-20827-4

27. Xu J, Casella JF, Pollard TD. Effect of capping protein, CapZ, on the length of actin filaments and mechanical properties of actin filament networks. Cell Motil Cytoskeleton. 1999;42: 73–81. doi:10.1002/(SICI)1097-0169(1999)42:1<73::AID-CM7>3.0.CO;2-Z

28. Wioland H, Guichard B, Senju Y, Myram S, Lappalainen P, Jégou A, et al. ADF/Cofilin Accelerates Actin Dynamics by Severing Filaments and Promoting Their Depolymerization at Both Ends. Current Biology. 2017;27: 1956–1967.e7. doi:10.1016/j.cub.2017.05.048

29. Raz-Ben Aroush D, Ofer N, Abu-Shah E, Allard J, Krichevsky O, Mogilner A, et al. Actin Turnover in Lamellipodial Fragments. Current Biology. 2017;27: 2963–2973.e14. doi:10.1016/j.cub.2017.08.066

30. Lai FPL, Szczodrak M, Block J, Faix J, Breitsprecher D, Mannherz HG, et al. Arp2/3 complex interactions and actin network turnover in lamellipodia. EMBO Journal. 2008;27: 982–992. doi:10.1038/emboj.2008.34

31. Gonzalez Rodriguez S, Wirshing ACE, Goodman AL, Goode BL. Cytosolic concentrations of actin binding proteins and the implications for in vivo F-actin turnover. Journal of Cell Biology. 2023;222. doi:10.1083/jcb.202306036

32. Sirotkin V, Berro J, Macmillan K, Zhao L, Pollard TD. Quantitative Analysis of the Mechanism of Endocytic Actin Patch Assembly and Disassembly in Fission Yeast. Schmid SL, editor. Mol Biol Cell. 2010;21: 2894–2904. doi:10.1091/mbc.e10-02-0157

33. Koestler SA, Rottner K, Lai F, Block J, Vinzenz M, Small JV. F- and G-Actin Concentrations in Lamellipodia of Moving Cells. Hotchin N, editor. PLoS One. 2009;4: e4810. doi:10.1371/journal.pone.0004810

34. Hackney DD, Clark PK. Steady state kinetics at high enzyme concentration. The myosin MgATPase. Journal of Biological Chemistry. 1985;260: 5505–5510. doi:10.1016/S0021-9258(18)89051-2

35. Abkarian M, Loiseau E, Massiera G. Continuous droplet interface crossing encapsulation (cDICE) for high throughput monodisperse vesicle design. Soft Matter. 2011;7: 4610. doi:10.1039/c1sm05239j

36. Spudich JA, Watt S. The regulation of rabbit skeletal muscle contraction. I. Biochemical studies of the interaction of the tropomyosin-troponin complex with actin and the proteolytic fragments of myosin. J Biol Chem. 1971;246: 4866–4871. doi:10.1016/s0021-9258(18)62016-2

37. Lappalainen P, Drubin DG. Cofilin promotes rapid actin filament turnover in vivo. Nature. 1997;388: 78–82. doi:10.1038/40418

38. D E Raeside. Monte Carlo principles and applications. Phys Med Biol. 1976;21: 181–197. doi:10.1088/0031-9155/21/2/001

39. Gillespie DT. Stochastic simulation of chemical kinetics. Annu Rev Phys Chem. 2007;58: 35–55. doi:10.1146/annurev.physchem.58.032806.104637

